# Unique and shared proteome responses of rice plants (*Oryza sativa*) to individual abiotic stresses

**DOI:** 10.1101/2022.10.19.512803

**Authors:** Fatemeh Habibpourmehraban, Brian J. Atwell, Paul A. Haynes

**Affiliations:** School of Natural Sciences, Faculty of Science and Engineering, Macquarie University, NSW 2109, Australia; Biomolecular Discovery Research Centre, Macquarie University, NSW 2109, Australia

**Keywords:** rice, abiotic stress, proteomics, shared responses, unique responses

## Abstract

Food safety of staple crops such as rice is of global concern and is at the top of the policy agenda worldwide. Abiotic stresses are one of the main limitations to optimizing yields for sustainability, food security and food safety. We analyzed proteome changes in *Oryza sativa* ssp. Nipponbare in response to three adverse abiotic treatments, including three levels of drought (mild, moderate, and severe), soil salinization, and non-optimal temperatures. All treatments had modest, negative effects on plant growth, enabling us to identify proteins that were common to all stresses, or unique to one. More than 75% of the total of differentially abundant proteins in response to abiotic stresses were specific to individual stresses, while fewer than 5% of stress-induced proteins were shared across all abiotic constraints. Stress-specific and non-specific stress-responsive proteins identified were categorized in terms of core biological processes, molecular functions, and cellular localization.

**Data Access:** All data have also been submitted to the PRIDE data repository, and will be available with project identifier PXD037280.

## 1. Introduction

Rice is one of the most valuable and important staple grains grown and consumed in our world, along with wheat and maize. According to the Food and Agriculture Organization data, world rice production has increased remarkably in recent years, with more than 80% used for food, but we still need more. Studies have predicted that world rice production will need to increase to 562.3 M metric tons by 2027, and rice consumption will increase at the same time to 459.5 metric tons. This is necessary for continuing to feed about half of the world population [1]. Therefore, the food security of staple crops such as rice is of global concern and has recently risen to the top of the policy agenda in many countries worldwide [2].

Hunger eradication is one of the main aims of the United Nations sustainable development goals. To feed up to 10 billion people in 2050, we need to improve sustainability, food security, and food safety, and make better use of food already produced. Some of the principal strategies for reducing food losses and waste can involve controlling environmentally unfavorable conditions, producing plant genotypes resistant to stresses, increasing the productivity and yield of staple plants like rice, and reusing or reprocessing surplus foods [3].

Based on currently available data, up to 30% of the main global food crops are lost annually due to plant biotic and abiotic stresses, with a considerable contribution from abiotic factors such as drought, salt, or temperature stresses [4]. Due to the long-term adverse effects of climate change, the frequency, intensity and duration of abiotic stresses are anticipated to increase in the coming years, posing serious threats to crop production and global food security [5]. Besides, abiotic stresses have been aggravated by the constant population growth, along with urbanization and the need to obtain a range of services from agriculture in addition to food production [6].

Plants are sessile organisms and cannot move away from stressful conditions, thus they must cope with all kinds of adverse external pressures via their intrinsic biological mechanisms [7]. Rice production in most of the cultivatable rice ecosystems, similarly to other food crops, has been progressively affected by various types of abiotic stresses, especially drought, salinity, heat, and cold [8].

In the post-genomics era, integrative omics can play a crucial role in studies of biochemical, physiological, and molecular analysis of plants in coping with stressful conditions. Large scale studies, which involve essentially developing an atlas of protein expression in plants under different abiotic stresses including salinity, drought, and temperature stress, have become increasingly important for discovering the potential key genes and proteins in different plant tissues [9]. Abiotic constraints reveal profound impacts on plant proteomes including alterations in protein relative abundance, cellular localization, post-transcriptional and post-translational modifications, protein interactions with other protein partners, and, finally protein biological functions [10]. Hence, the role of proteins in plant stress response is crucial since proteins are directly involved in shaping novel phenotypes by adjustment of physiological traits to cope with altered environments, and proteomic analysis technologies can be used to monitor and characterize differences in protein expression profiles in rice plants.

In this study, a low land rice genotype, Nipponbare, with low tolerance to abiotic stresses, especially water deficiency [11, 12], was exposed to single abiotic stresses separately, including different drought stress levels (mild, moderate, and severe), salt stress and temperature stress, to analyze the shared and specific stress proteome responses. Protein abundances were investigated using label-free shotgun proteomics and subsequent data analysis revealed the role of protein biological function and cellular localization on protein stress response specificity.

## 2. Results

### 2.1 Label-free shotgun proteomics data analysis

Seedlings of lowland Nipponbare rice plants, which is considered to be an abiotic stress-sensitive genotype [11], were grown for four weeks followed by exposure to three separate abiotic stresses; drought, salt, and temperature stress treatments. Plants were exposed to three different levels of water deficit by reducing the soil field capacity of watering (FC) to 70%, 50%, and 30%, referred to hereafter as mild, moderate and severe drought. The concentration of sodium chloride in the soil was increased to 50mM for another set of rice plants, creating salt stress, and the last group of plants was treated with unfavorable temperature conditions by increasing the day-time temperature to 33°C and decreasing the night-time temperature to 18°C. Leaf samples from stressed and control plants were taken for proteomics analysis, to characterize the shared and unique proteome responses of the Nipponbare genotype to individual abiotic stresses.

Table 1 shows the number of reproducibly identified proteins and peptides from the analysis of each set of plants including control and all five stress treatments. A non-redundant total of 3997 proteins were identified across all treatments and groups when all six sample data sets were combined. The highest number of reproducibly identified proteins was 1732 in control conditions, while the lowest number was 1013 reproducibly identified proteins in moderate drought stress.

**Table 1.** Summary of the protein and peptide quantification data identified under individual abiotic stresses in Nipponbare (FDR= False Discovery Rate).

| Stress Treatment | Reproducibly identified proteins | Reproducibly identified peptides | Protein FDR % | Peptide FDR % |
| --- | --- | --- | --- | --- |
| Control | 1732 | 93277 | 0.29 | 0.07 |
| Mild Drought | 1256 | 67648 | 0.48 | 0.14 |
| Moderate Drought | 1013 | 56448 | 0.20 | 0.04 |
| Severe Drought | 1182 | 64351 | 0.34 | 0.11 |
| Salt | 1587 | 130290 | 0.95 | 0.31 |
| Temperature | 1426 | 81710 | 0.21 | 0.05 |

Table 2 shows the number of differentially expressed proteins (DEPs) identified as being statistically significantly changed in abundance for each of the stress treatments. Increasing the drought stress severity from mild to severe increased the number of differentially abundant proteins. However, the increase in DEPs between mild and moderate drought was much greater than that which occurred between moderate and severe drought. Both moderate and severe drought cause more than 20% of the reproducibly identified proteins to be changed in abundance, which represents a greater systemwide disturbance than that seen for the other stress conditions. Details of all proteins and DEPs identified are provided in supplementary data file S1.

**Table 2.** Differentially expressed proteins (DEPs) under individual abiotic stresses in Nipponbare.

| Stress Treatment | DEPs | Increased in abundance | Decreased in abundance | % changed (of total number identified) |
| --- | --- | --- | --- | --- |
| Mild Drought | 267 | 66 | 201 | 13.9 |
| Moderate Drought | 405 | 80 | 325 | 21.8 |
| Severe Drought | 420 | 119 | 301 | 21.7 |
| Salt | 307 | 68 | 239 | 15.0 |
| Temperature | 185 | 56 | 129 | 9.3 |

### 2.2 Analyzing the effect of divergent drought stress levels on leaf proteome quantification

A total of 1092 proteins were changed in abundance in response to various drought stress levels in Nipponbare plants, but less than 10% of these (106) were changed in abundance in response to all three drought conditions, indicative of a shared stress response (Figure 1). Moreover, the greatest number of commonly altered proteins was observed between moderate and severe drought (31 proteins increased in abundance and 83 proteins decreased in abundance). The 52% overlap between differentially abundant proteins at these two drought stress levels illustrates that Nipponbare plants sensed and responded to moderate and severe drought stresses quite similarly. Figure 1 also demonstrates the number of unique and shared differentially abundant proteins seen in diverse drought conditions, with the highest number of distinct proteins increased in abundance in response to severe drought with 51 proteins, while in comparison the greatest number of unique proteins decreased in abundance was 127 proteins altered in response to moderate drought. The pie charts in Figure 1 show that the total number of differentially abundant proteins identified uniquely in response to a single stress was approximately 10% higher than the total number of differentially abundant proteins identified in more than one condition.

**Figure 1.**
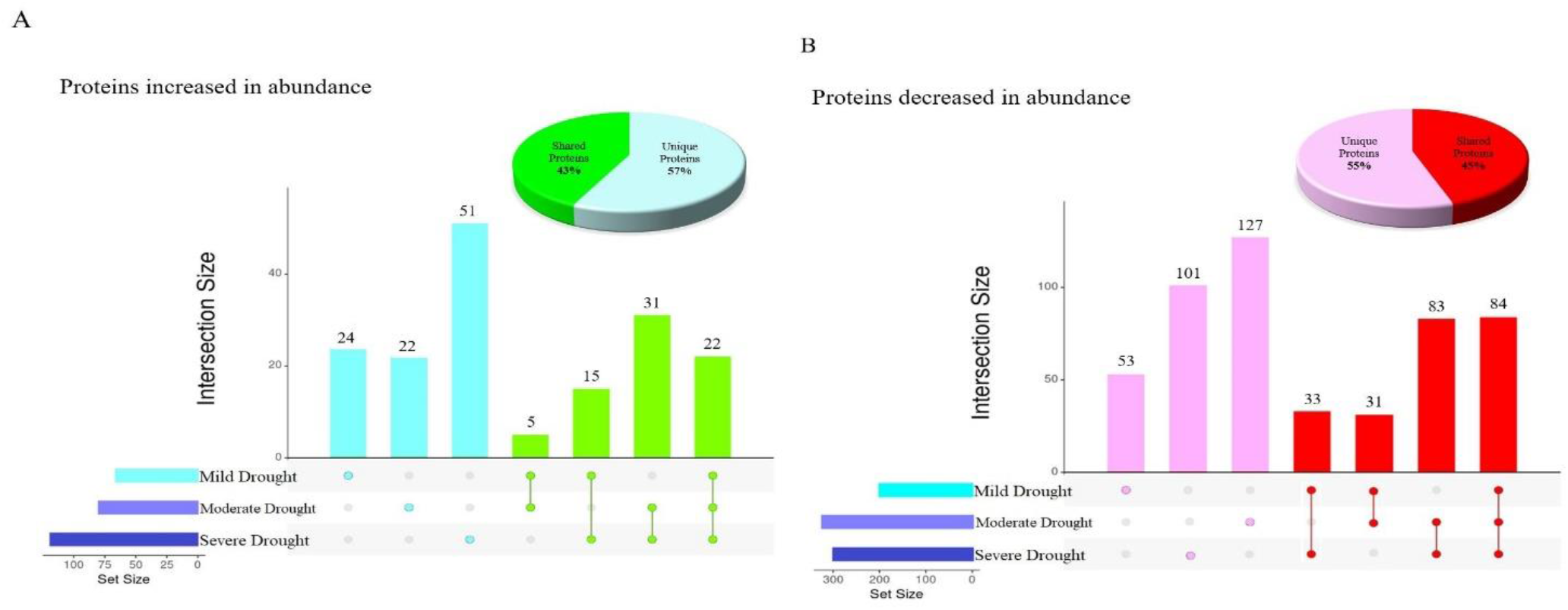
Overlap between the proteins differentially expressed under various levels of drought stress in Nipponbare. (A) Upset plot [13] indicating the overlap between proteins significantly increased in abundance and Pie chart representing the proportion of unique and shared proteins significantly increased in abundance (B) Upset plot indicating the overlap between proteins significantly decreased in abundance and pie chart representing the proportion of unique and shared proteins significantly decreased in abundance.

Almost all of the proteins that were increased or decreased in abundance in more than one drought stress condition were similarly increased or decreased in all stress conditions where they were observed, except for two proteins with very unusual patterns of abundance (Table 3). C2H2-type domain-containing protein (Q8GS72) was decreased in abundance in response to severe drought, but increased in abundance in response to mild drought stress. Putative glycine-rich protein (Q6Z142) was decreased in abundance in moderate drought, but increased in abundance at mild drought stress level. C2H2 zinc finger proteins have been shown to participate as a pivotal regulator of Reactive Oxygen Species (ROS) signaling in the signal transduction of water stress [14], and oxidative stress, in rice [15]. It has been demonstrated previously that repressing the expression of zinc finger proteins under drought stress was linked to enhancing tolerance to drought stress through a complex regulatory network in plants [16]. This suggests that Nipponbare plants sensed the severe drought condition as stress and responded to it by decreasing the abundance of C2H2-type domain-containing protein.

**Table 3.** Protein abundance (fold-change) under various drought stress conditions for two proteins with unusual patterns of changes in abundance

| Protein ID | Q8GS72 | Q6Z142 |
| --- | --- | --- |
| Protein name | C<sub>2</sub>H<sub>2</sub>-type domain-containing protein | Putative glycine-rich protein |
| Severe Drought | -1.73 | Unchanged |
| Moderate Drought | Unchanged | -5.99 |
| Mild Drought | 1.17 | 3.26 |

Glycine-rich RNA-binding proteins (GRPs), such as putative glycine-rich protein (Q6Z142), have been implicated in the responses of plants to environmental stresses including dehydration, and it has been determined that activation of GRP-related genes has a negative impact on plant response to water deficiency stress [17]. This agrees with our finding that putative glycine-rich protein is increased in case of mild drought stress conditions, but then decreased in abundance under moderate drought stress.

### 2.3 Shared functional proteome response under three drought stress levels

Proteomic investigation of plants grown under drought stress conditions identified differentially abundant proteins in many different functional classifications, which provide insights into water deficit proteome response mechanisms. Changes in abundance of proteins in many different functional categories reflects the diversity and versatility of plant responses when exposed to different levels of water deficit.

Figure 2A displays the gene ontology (GO) analysis of the 106 proteins that were changed in abundance under all three drought stress conditions. This reveals that the proteins commonly changed in response to drought stress were mostly related to photosynthesis, translation, and transport functions. Proteins in the photosynthesis functional category were mostly increased in abundance while the majority of proteins in the translation and transport categories were decreased in abundance. Growth was the only functional category in which all shared proteins were decreased in abundance.

**Figure 2.**
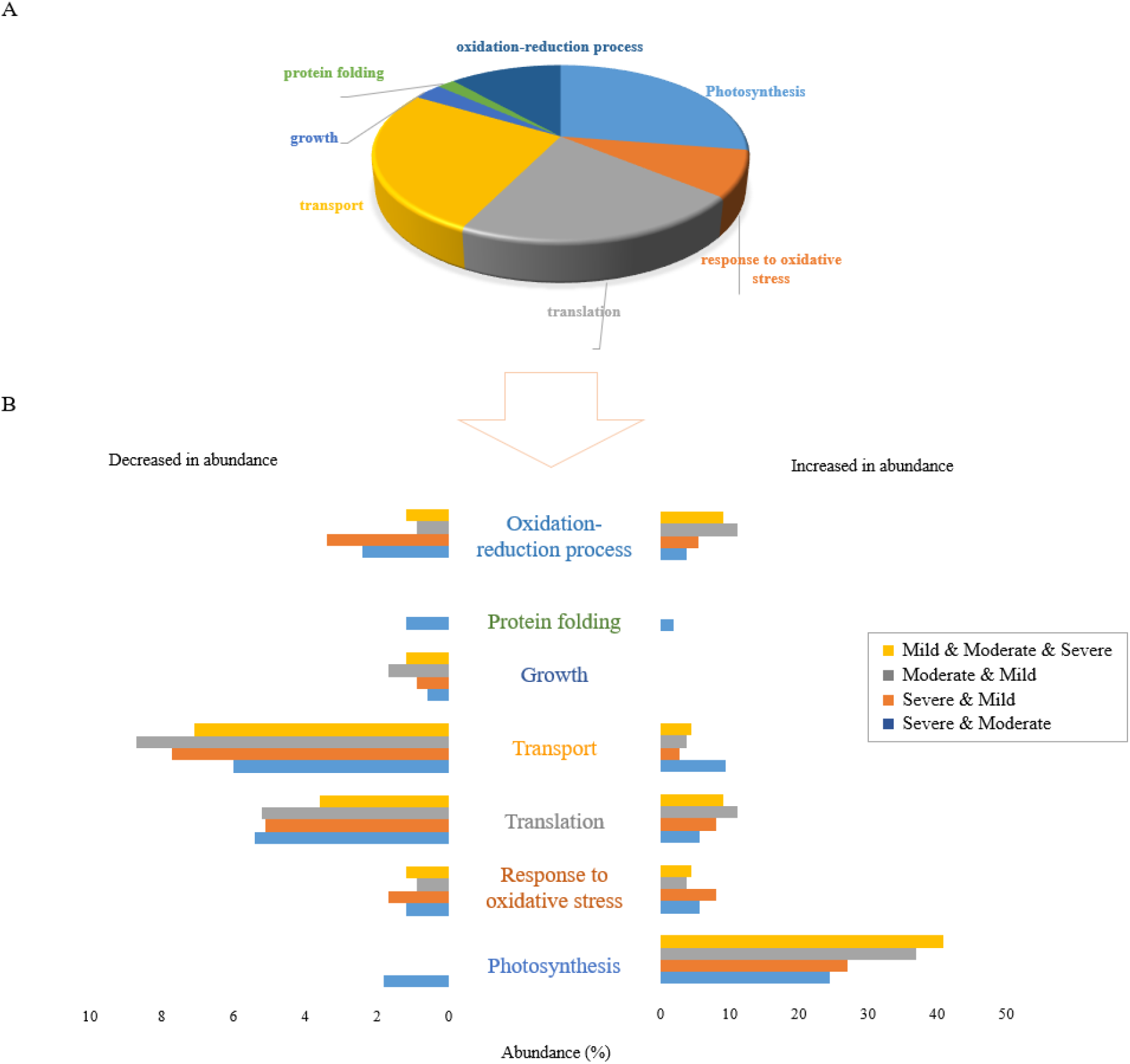
Functional classification of 106 differentially accumulated proteins in response to all 3 drought stress levels. A) The pie chart shows the percentage of different biological functions of common proteins significantly changed in abundance in response to all three drought stress levels. B) The bar graph shows the percentage abundance of proteins in 7 functional categories that were significantly altered in response to drought stress at divergent levels. Distinct colours represent proteins significantly changed in common between stresses: severe and moderate drought (blue), severe and mild drought (orange), moderate and mild drought (grey), and overlapped between all three drought stress levels (yellow).

The proportional abundance of proteins increased and decreased commonly between combinations of two or three drought stress levels is represented in Figure 2B. Clearly, the greatest effect of drought stress occurred in proteins increased in abundance in the photosynthesis functional category. In terms of proteins decreased in abundance in response to combinations of stresses, the highest proportion of proteins was those related to the transport function, with proteins in the translation functional category the next most abundant. These results reflect the generalised effects of drought stress on Nipponbare plants with respect to stress level.

### 2.4 Stressor-specific proteome response under individual drought stress levels

The results of proteomic studies can reflect different stress-coping strategies depending on the given stress treatment. Analysis of the GO functional categorization of 378 proteins specifically expressed under each drought stress condition is shown in Figure 3 (152 proteins in severe drought, 149 proteins in moderate drought and 77 proteins in mild drought). Similar to the shared proteins, photosynthesis, transport, and translation were the main biological functions affected by individual drought stresses. Proteins in the growth functional category were all decreased in abundance (Figure 3A). One such protein that was significantly decreased in abundance only in plants subjected to severe drought stress was putative acyl-CoA dehydrogenase (Q6ZDX3). Increasing acyl-CoA dehydrogenase accumulation under stress treatment has been reported to be a molecular response of plants which enables them to grow more rapidly, so decreasing the protein abundance under severe drought reflects the negative impact on plant growth of severe drought conditions. Except for growth, all six other biological functions responded to severe drought stress level by both increasing and decreasing the abundance of related proteins (Figure 3B).

**Figure 3.**
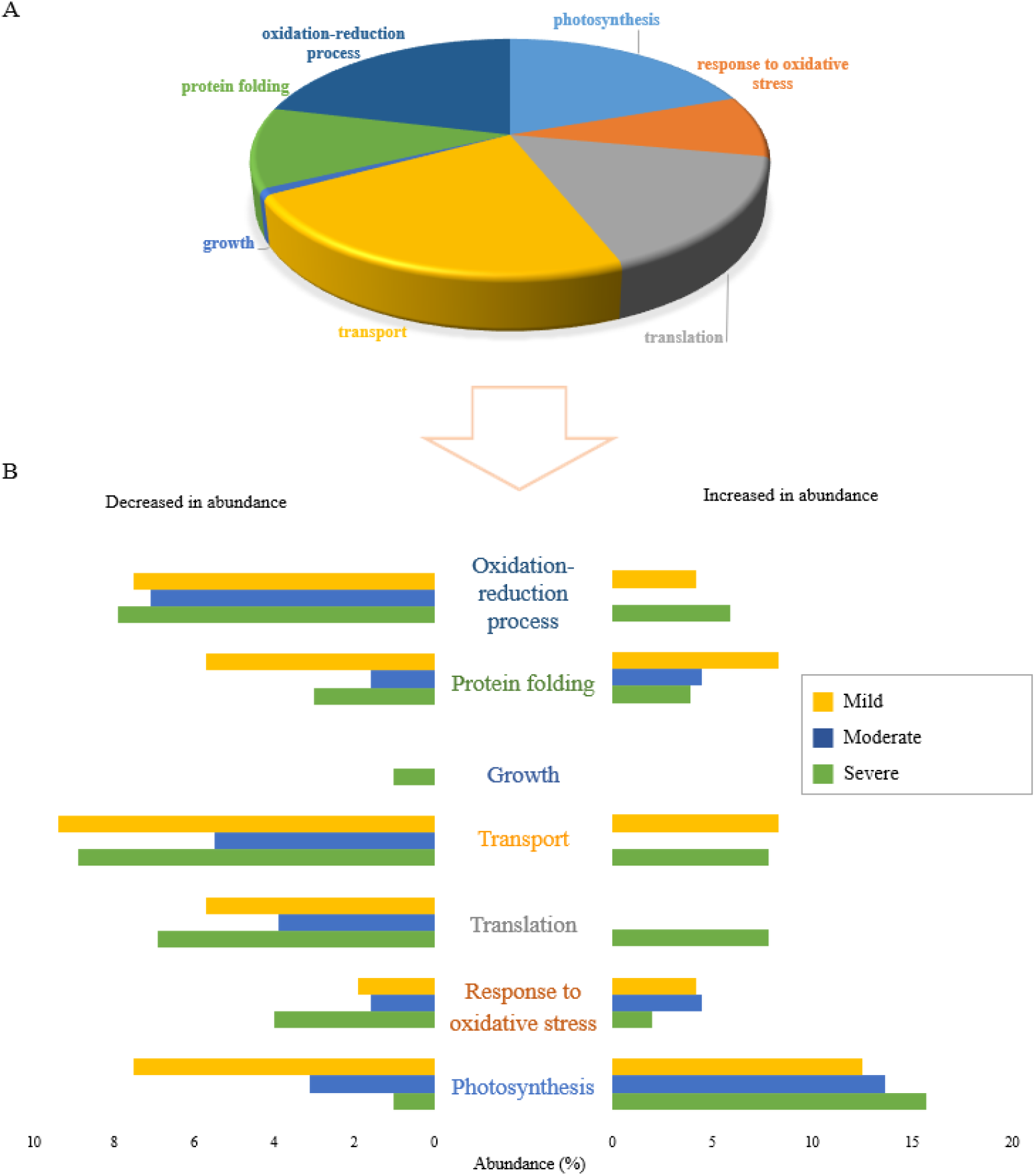
Functional classification of proteins differentially accumulated in response to drought stress levels, specific to each of severe, moderate, and mild drought stresses. A) The pie chart shows the percentage of biological functions of 378 proteins significantly changed uniquely under different drought stress levels. B) The bar graph shows the abundance of proteins in percentage in 7 functional categories that are uniquely altered in response to drought stress levels including severe, moderate, and mild drought treatments. Distinct colours represent various levels of field capacity in drought stress: severe drought specific (green), moderate drought specific (blue) and mild drought specific (yellow).

Analysis of biological functional categories of proteins changed in response to drought stress, either uniquely or shared between different conditions, indicated that severe drought stress appeared to cause the most obvious changes in the expressed proteome of Nipponbare plants, irrespective of whether proteins decreased or increased in abundance.

### 2.5 Proteome quantification comparison under various abiotic stresses

Different environmentally unfavourable conditions lead to both common and unique molecular processes in plants. The specific proteins and protein expression patterns under various stress treatments reflect the evolution of diverse plant tolerance mechanisms. Moreover, despite various stress-independent commonalities when plants are exposed to a particular abiotic stress, stress-specific signatures can also be detected in plants.

Prior to examining the common proteome response when plants were exposed to drought, salt, and temperature stresses, we needed to choose the best candidate for subsequent comparisons from among the different drought stress levels. Considering that the overlapped proteins differentially expressed under all three different drought stress levels gave us a small number of proteins (Figure 1) for further comparative proteome analysis with salt and temperature treatments, we decided to focus on common proteins expressed under two drought stress conditions. Hence, common proteins differentially altered in abundance in both moderate and severe drought stresses (53 proteins increased in abundance and 167 proteins decreased in abundance) were considered as proteins affected by drought stress in Nipponbare plants, and included in subsequent comparative analyses with salt and temperature stress.

Figure 4 shows that approximately 75% of 551 total differentially expressed proteins were stressor-specific proteins and only less than 5% (24 proteins) belonged to shared responses among all abiotic stresses. Approximately half of the proteins changed in abundance by only one stress treatment were those induced in response to salt stress treatment, with 156 proteins decreased and 46 proteins increased in abundance.

**Figure 4.**
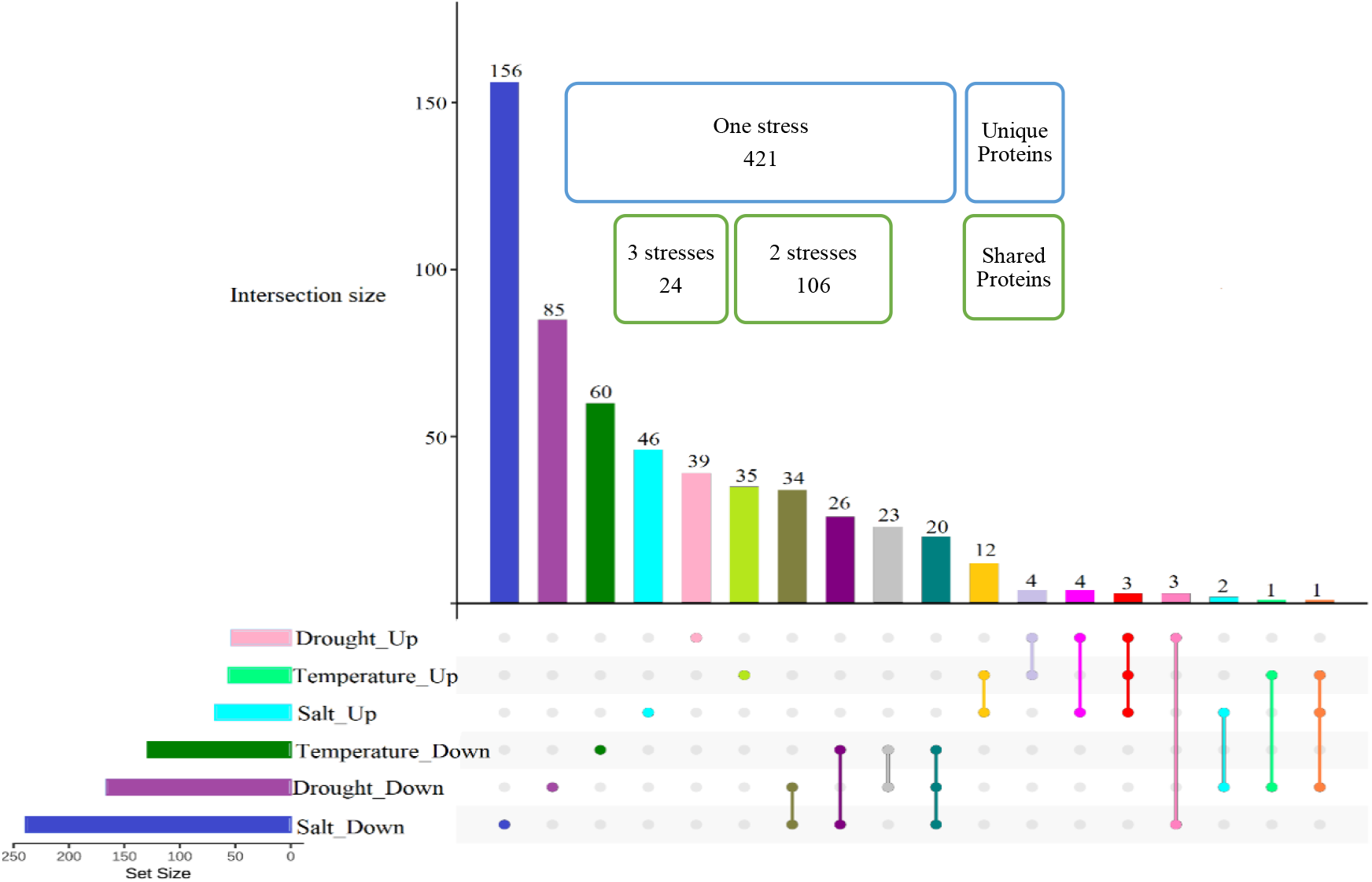
Number of proteins significantly differentially abundant uniquely and in common in response to salt, drought, and temperature. The Upset plot shows the proteins significantly differentially expressed under individual abiotic stresses, and shared with other stresses. The two highlighted groups show the number of proteins specific to each abiotic stress (unique proteins), and the number of proteins shared among abiotic stresses (shared proteins).

### 2.6 Proteins induced in common under abiotic stresses

Figure 4 and Table 4 show that 24 proteins were changed in abundance in response to all three abiotic stresses, with 20 proteins decreased in abundance in response to all three stresses and 3 increased in abundance in response to all three stresses; Epimerase domain-containing protein (Q2QSR7), L-ascorbate peroxidase 8 (Q69SV0), and Phosphoglucomutase (Q9AUQ4). The only protein with changes in abundance occurring in contrasting directions in response to different stresses was Peptidyl-prolyl cis-trans isomerase (Q69WA8), with a similar increase in abundance of 1.99-fold change under both temperature and salt stresses, and significant decrease in abundance under drought conditions (Table 4). Interestingly, the level of protein abundance for Epimerase domain-containing protein (Q2QSR7) and L-ascorbate peroxidase 8 (Q69SV0) proteins increased further with decreasing the level of soil water, while the trend for phosphoglucomutase, along with that of (Q69WA8), was in the opposite direction, with protein abundance decreasing when the drought stress was increased from moderate to severe. The unusual behavior of these proteins in Nipponbare plants, where they are altered in abundance in subtly different ways in response to different individual abiotic stresses, makes them attractive candidates for future detailed studies

**Table 4.** Heatmap Table of the relative expression level of DEPs involved in three abiotic stresses including severe drought, moderate drought, salt, and temperature stresses in Nipponbare plants.

| Protein name | Protein ID | Temperature | Salt | Severe Drought | Moderate Drought |
| --- | --- | --- | --- | --- | --- |
| Epimerase domain-containing | Q2QSR7 | 1.64 | 1.67 | 1.81 | 1.77 |
| L-ascorbate peroxidase 8 | Q69SV0 | 1.41 | 1.49 | 1.61 | 1.38 |
| Phosphoglucosmutase | Q9AUQ4 | 1.36 | 1.48 | 1.75 | 2.26 |
| Glucan endo-1,3-beta-D-glucosidase | A0A0P0XDW1 | -3.77 | -7.14 | -7.25 | -3.83 |
| Beta-glucosidase 31 | B7F7K7 | -4.63 | -7.76 | -4.51 | -3.86 |
| Aspergillus nuclease S(1) | B9FCW0 | -9.44 | -5.64 | -5.49 | -4.69 |
| Aldo_ket_red domain-containing | Q0JCV5 | -4.85 | -7.09 | -7.19 | -3.47 |
| W2 domain-containing protein | Q2R678 | -2.22 | -1.86 | -1.52 | -1.88 |
| DEK C-terminal domain-containing | Q5JKH1 | -2.87 | -2.32 | -2.06 | -2.40 |
| 5'-3' exoribonuclease | Q5N739 | -7.25 | -2.11 | -4.25 | -6.03 |
| Ribokinase (RK) | Q5SN59 | -2.00 | -2.09 | -6.11 | -3.90 |
| Abhydrolase_3 domain-containing | Q5Z4C9 | -3.47 | -2.45 | -2.67 | -3.48 |
| Hydrolase_4 domain-containing | Q5ZC21 | -3.90 | -3.74 | -3.79 | -3.24 |
| Transket_pyr domain-containing | Q69LD2 | -2.00 | -2.74 | -2.01 | -3.22 |
| FAD-binding FR-type domain-containing | Q69LJ7 | -3.85 | -6.44 | -3.73 | -3.18 |
| Endoglucanase | Q69NF3 | -3.32 | -2.84 | -3.15 | -3.39 |
| Glycin-rich RNA-binding protein | Q6ASX7 | -2.09 | -2.46 | -33.80 | -28.88 |
| Transcription factor GTE8 | Q6K5G2 | -4.83 | -5.38 | -4.67 | -3.99 |
| F-box domain-containing | Q6Z4S6 | -3.68 | -3.31 | -2.19 | -3.86 |
| Calcium-dependent protein kinase 8 | Q75GE8 | -4.12 | -5.97 | -6.11 | -5.22 |
| Fasciclin-like arabinogalactan protein 15 | Q7XIM4 | -3.12 | -10.27 | -5.97 | -5.11 |
| Calmodulin-binding protein 60 C | Q7XRM0 | -4.54 | -11.56 | -6.73 | -5.75 |
| t-SNARE coiled-coil homology domain-containing | Q8H5R6 | -5.42 | -5.18 | -5.25 | -4.49 |
| Peptidyl-prolyl cis-trans isomerase | Q69WA8 | 1.99 | 1.99 | -4.54 | -3.88 |

### 2.6 Proteins induced in common under abiotic stresses

Figure 4 and Table 4 show that 24 proteins were changed in abundance in response to all three abiotic stresses, with 20 proteins decreased in abundance in response to all three stresses and 3 increased in abundance in response to all three stresses; Epimerase domain-containing protein (Q2QSR7), L-ascorbate peroxidase 8 (Q69SV0), and Phosphoglucomutase (Q9AUQ4). The only protein with changes in abundance occurring in contrasting directions in response to different stresses was Peptidyl-prolyl cis-trans isomerase (Q69WA8), with a similar increase in abundance of 1.99-fold change under both temperature and salt stresses, and significant decrease in abundance under drought conditions (Table 4). Interestingly, the level of protein abundance for Epimerase domain-containing protein (Q2QSR7) and L-ascorbate peroxidase 8 (Q69SV0) proteins increased further with decreasing the level of soil water, while the trend for phosphoglucomutase, along with that of (Q69WA8), was in the opposite direction, with protein abundance decreasing when the drought stress was increased from moderate to severe. The unusual behaviour of these proteins in Nipponbare plants, where they are altered in abundance in subtly different ways in response to different individual abiotic stresses, makes them attractive candidates for future detailed studies.

Among the 20 proteins that were reduced in abundance in response to all three abiotic stresses in all plants, Glycine-rich RNA-binding protein (Q6ASX7) displayed the largest variation in expression between drought stress and other abiotic stresses. Other examples of variation in expression level under different stresses were Fasciclin-like arabinogalactan protein 15 (Q7XIM4) and Calmodulin-binding protein 60 C (Q7XRM0) which decreased in abundance under all abiotic stresses, but with a much greater change in abundance in response to salinity, and Putative 5’-3’ exoribonuclease (Q5N739) which decreased much more under temperature stress in comparison to the other two stresses. Investigating the metabolic activities of each of these proteins could help us to understand the mechanistic reasons for differences in abundance change of each in response to different individual abiotic stress.

A total of 106 proteins were changed in abundance under any combination of two stress conditions (Figure 4). This included both similar and antithetic reactions of plants to different stresses; 6 of these showed abundance changes in opposite direction between drought, and each of salt or temperature stresses. Specifically, UDP-glucose 6-dehydrogenase 3 (Q9AUV6) was changed in abundance in response to drought and temperature stresses, however, its abundance decreased under drought stress and increased in response to temperature extreme. Notably, five proteins responded to both drought and salt stresses in common but in opposite manner. Two of them were chloroplastic proteins, namely Glutamate-1-semialdehyde 2,1-aminomutase (Q6YZE2) and Uroporphyrinogen decarboxylase 2 (Q10LR9), which increased in abundance under salt stress but decreased in abundance in case of water deficiency. Three other proteins with different cellular localizations showed an opposite pattern by decreasing in abundance in response to salinity and increasing in abundance in response to drought stress, including Peroxiredoxin Q (P0C5D5), actin (Q67G20), and aquaporin PIP2-7 (Q651D5).

### 2.7 Determining the molecular functions of stress-specific and non-specific proteins under individual abiotic stresses

Analysis of proteins at the organelle level is a useful approach for understanding cell behavior under abiotic stress conditions. Abiotic stress alters interactions between organelles in plant cells, and this subsequently changes the regulation and secretion of proteins in cellular organelles and compartments. Therefore, to better understand the biological function of stress-specific and shared differentially abundant proteins under unfavourable conditions, we focused on the molecular functions of major subcellular organelles where functions are typically affected under abiotic stress, including the nucleus, cytosol, plasma membrane, cell wall, mitochondria, cytosol, vacuole, chloroplast and endoplasmic reticulum (Figure 5).

**Figure 5.**
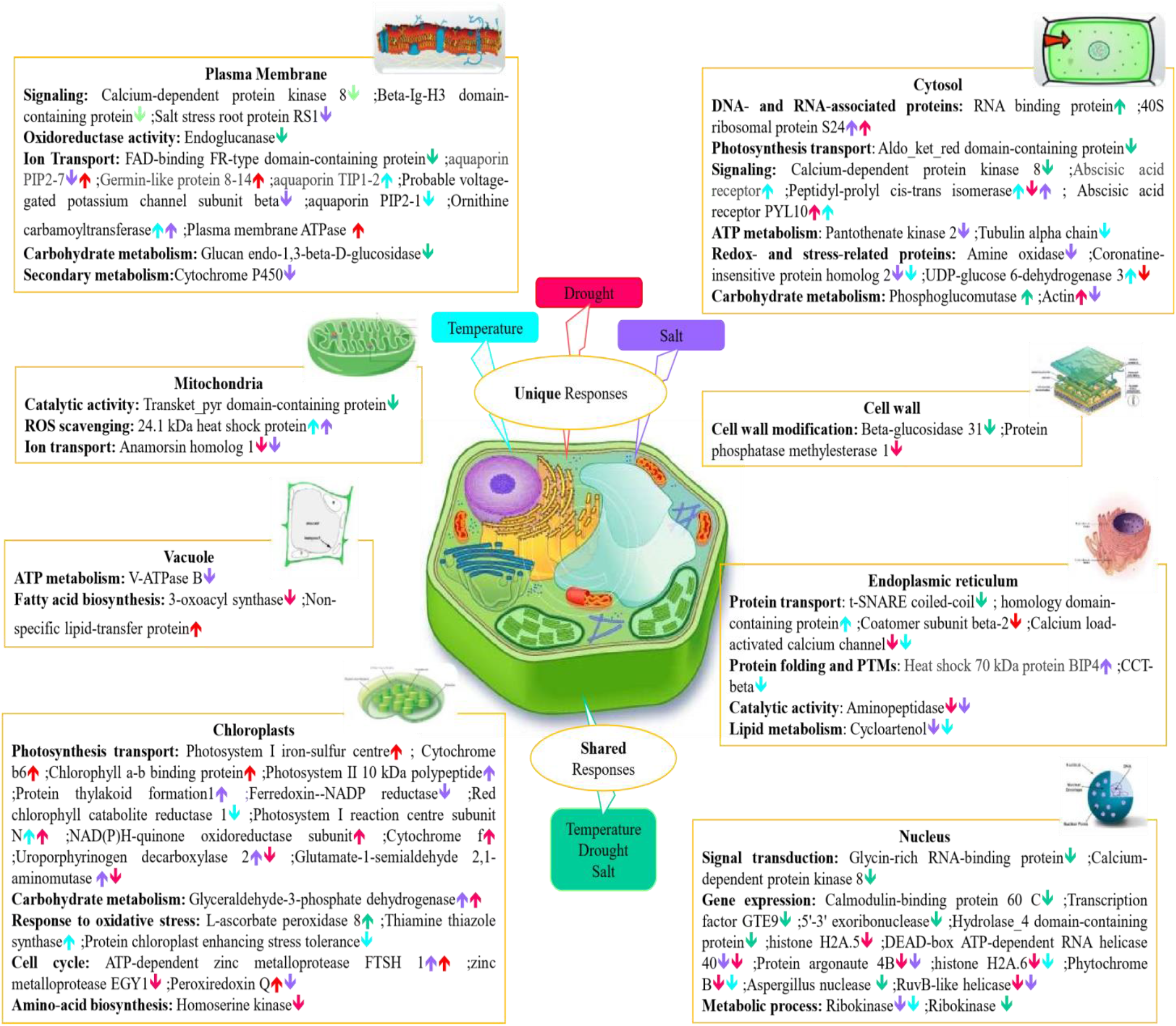
A schematic summary of the major stress-responsive proteins identified in this study, presented in context of their individual plant cell compartments. Differentially abundant proteins under drought, salt, and temperature stress are highlighted with distinct colours including salt unique proteins (purple), temperature unique proteins (blue), drought unique proteins (red), and shared proteins among all abiotic stresses (green). Upward and downward arrows indicate proteins increased and decreased in abundance.

To withstand environmental stresses, plants have evolved interconnected regulatory pathways that enable them to respond and adapt to their environments promptly. Plant stress proteomics is a dynamic discipline aimed at the study of plant acclimation or tolerance regulations and mechanisms exposed to stress. However, the differential expression proteomics approach itself, based on protein identification and quantitation, can sometimes not give sufficient information on protein function since a certain protein can be responsible for diverse functions depending on its subcellular localization.

Proteomic analyses of Nipponbare plant leaves under various abiotic stress conditions can also be employed for the identification of molecular and cellular mechanisms that are specific to certain abiotic stresses, or shared between two or more abiotic stress treatments. On the other hand, apart from the unique protein dynamic change mediated by different stress conditions, there can be various points of crosstalk between stress signaling pathways. Identification of crosstalk between signaling pathways has been crucial in strengthening our understanding of how plants regulate their responses to a particular stress condition. For example, aquaporin family proteins that are responsible for ion transport in the plasma membrane responded both negatively and positively to stress in this study. Specifically, aquaporin PIP 2-7 protein was increased in abundance under temperature and salt stress but decreased in abundance when coping with drought stress.

Chloroplasts play a key role in plant response to stress treatment, by adjustment of photosynthesis as a biological function. All individual abiotic stresses affected chloroplasts both negatively and positively, either uniquely or in common. For example, L-ascorbate peroxidase 8, which is a chloroplastic protein responsible for oxidative stress response, was reduced in abundance in response to all abiotic stresses. Some proteins related to different molecular mechanisms in chloroplasts showed opposing changes in abundance under drought and salt stresses, such as Peroxiredoxin, Uroporphyrinogen decarboxylase, and Glutamate-1-semialdehyde 2,1-aminomutase.

Abundance of DEPs in the nucleus were mainly decreased in response to abiotic stresses in both shared and stressor-specific protein categories. For instance, nuclear proteins related to major molecular functions such as gene dynamic change, signal transduction, and metabolic process, were decreased in abundance under all abiotic stresses. One example of differential effects of various abiotic stresses on plant cell organelle functions in our study was related to the heat shock protein family (HSPs). HSPs are known as one of the major stress responses mostly involved in response to temperature stress. Although 24.1 kDa heat shock protein, which is localized in mitochondria, was increased in abundance in response to both temperature and salt stresses, another member of this family, Heat shock 70 kDa protein BIP4, which occurs in the endoplasmic reticulum (ER), was increased in abundance only under salt stress.

The cytosol plays a significant role in plant function under stresses, not only by itself but also through interaction among different subcellular organelles. RNA binding protein and Phosphoglucomutase were two cytosol-localized proteins that were accumulated in common among all abiotic stresses. However, Aldo_ket_red domain-containing protein and Calcium-dependent kinase, which are related to photosynthesis and transport functions, respectively, decreased in abundance commonly under all individual abiotic stresses. Peptidyl-prolyl cis-trans isomerase, which performs a signaling molecular function in the cytosol, was detected as the only protein that increased in abundance under drought stress but decreased in abundance under both salt and temperature stress treatments.

### 2.8 Parallel Reaction Monitoring (PRM) validation

A series of PRM experiments were performed to validate the results from label-free shotgun proteomics analysis of Nipponbare plants under various individual abiotic stresses. The PRM results indicated that the differential changes in abundance of four selected proteins measured by PRM, as shown in Figure 6, agreed with the label-free shotgun proteomics results.

**Figure 6.**
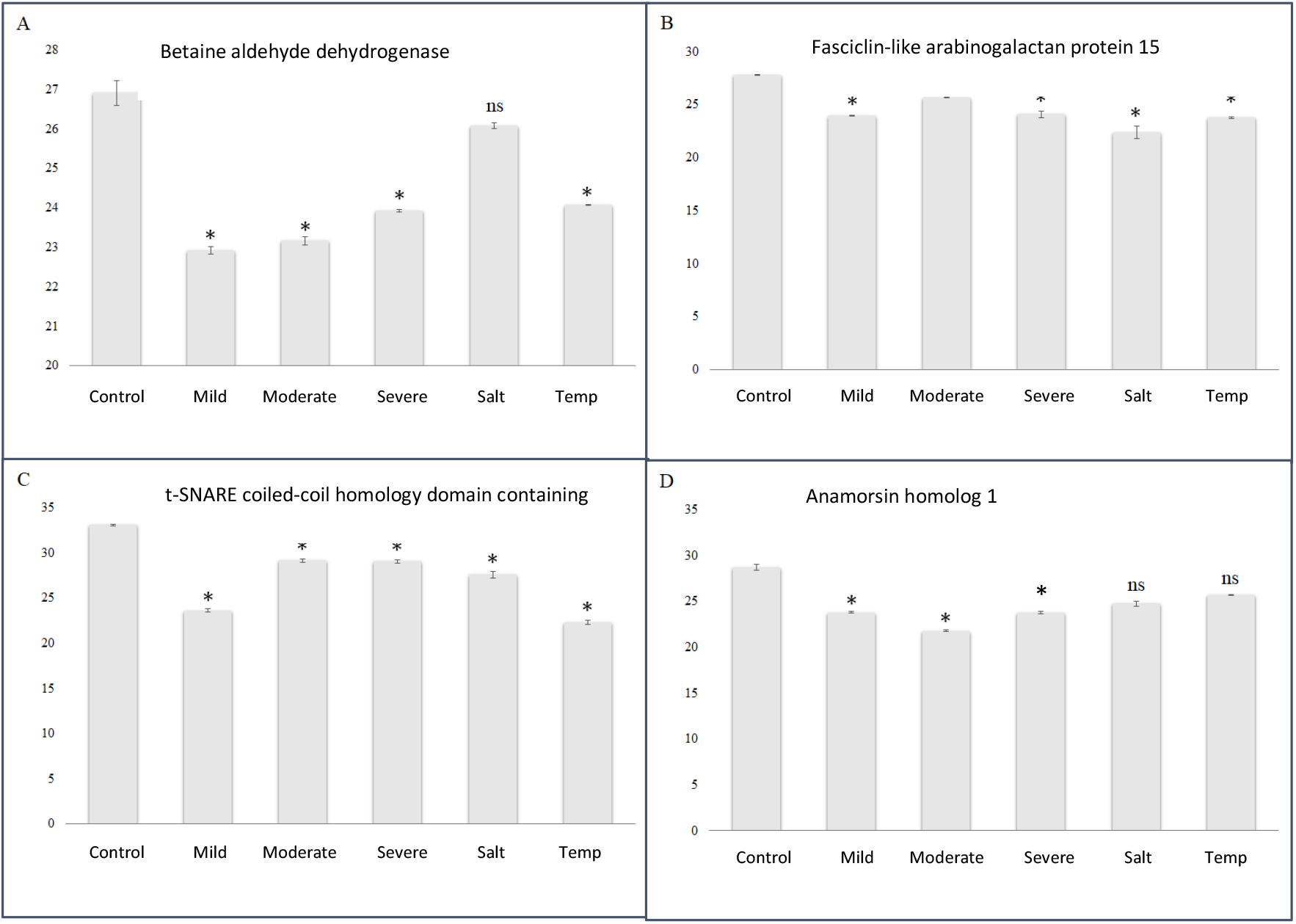
PRM validation of 4 proteins changed in abundance in response to abiotic stresses in Nipponbare plants. Bars illustrate the relative abundance of proteins under control conditions, followed by mild, moderate and severe drought stresses, salt stress and temperature stress treatments. An asterisk (*) indicates a statistically significant difference between the control and stress condition while (ns) indicates no statistically significant change between control and stress condition, according to a Student t-test (p-value < 0.05).

These included Betaine aldehyde dehydrogenase (O24174), Fasciclin-like arabinogalactan protein 15 (Q7XIM4), t-SNARE coiled-coil homology domain-containing (Q8H5R6), and Anamorsin homolog 1 (Q7XQ97). Betaine aldehyde dehydrogenase protein was significantly reduced in abundance under drought and temperature stresses, and was reduced in expression slightly after salt treatment but not to a statistically significant level (Figure 6 A). The Fasciclin-like arabinogalactan protein 15 (Figure 6B) and t-SNARE coiled-coil homology domain-containing protein (Figure 6C) both responded similarly to all abiotic stresses by decreasing in abundance significantly. Anamorsin homolog 1 responded to drought stresses by decreasing in abundance significantly, while under salt and temperature stress it was reduced in abundance, but not to a statistically significant level (Figure 6D). Table 5 shows for comparison purposes the reported fold-change of each of the four proteins from the label free quantitative proteomics data analysis.

**Table 5.**
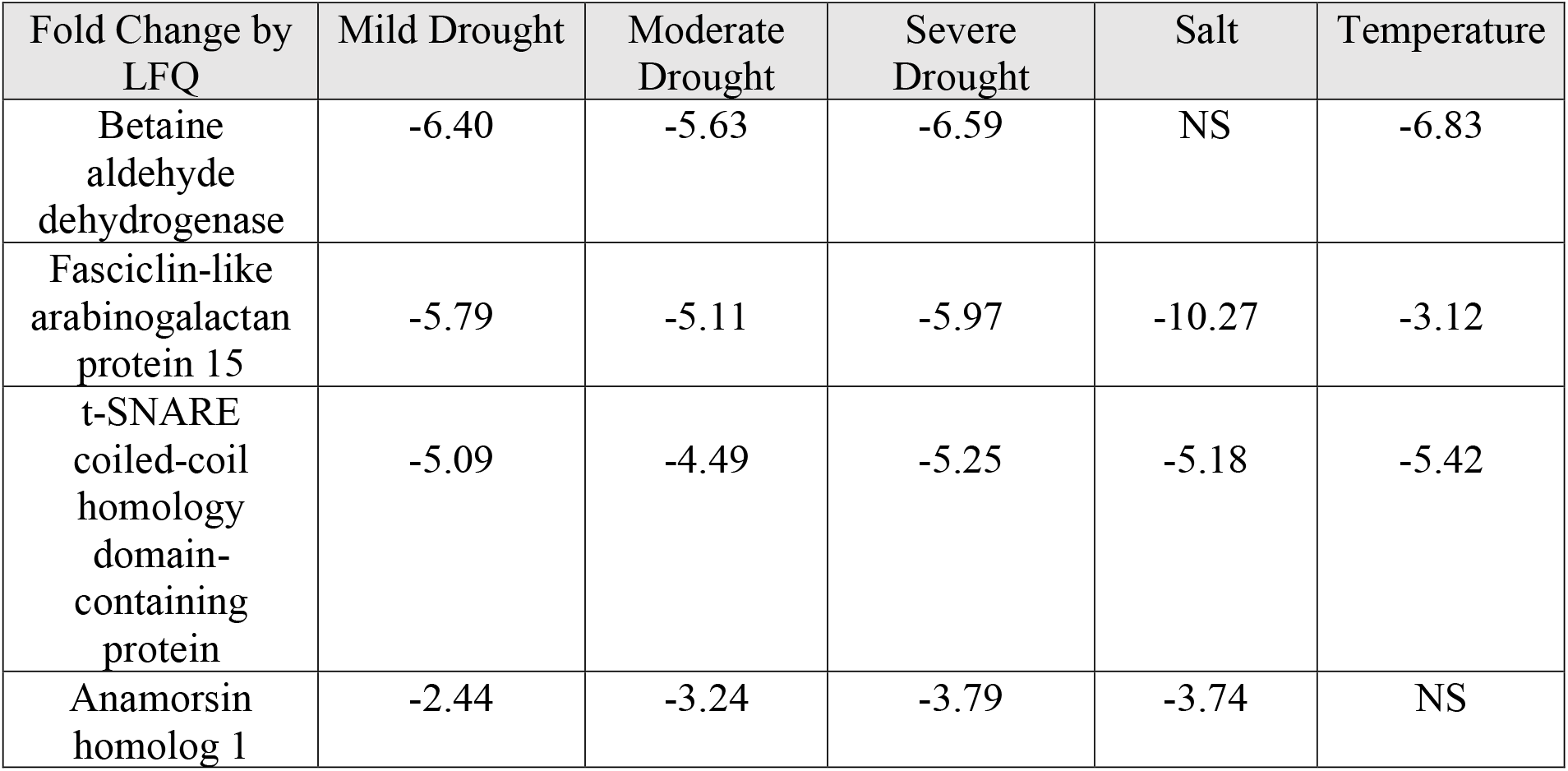
Fold change of proteins from LFQ experiment under different abiotic stresses in comparison with control plants. (NS) indicates no statistically significant change between control and stress condition, according to a Student t-test (p-value < 0.05).

| Fold Change by LFQ | Mild Drought | Moderate Drought | Severe Drought | Salt | Temperature |
| --- | --- | --- | --- | --- | --- |
| Betaine aldehyde dehydrogenase | -6.40 | -5.63 | -6.59 | NS | -6.83 |
| Fasciclin-like arabinogalactan protein 15 | -5.79 | -5.11 | -5.97 | -10.27 | -3.12 |
| t-SNARE coiled-coil homology domain-containing protein | -5.09 | -4.49 | -5.25 | -5.18 | -5.42 |
| Anamorsin homolog 1 | -2.44 | -3.24 | -3.79 | -3.74 | NS |

## 3. Discussion

Plant responses to various stressful environmental conditions can be considered as two main categories including shared responses and responses which are unique to specific plants or specific stressors. The sensitivity or tolerance of each plant to a variety of stress conditions can be differentiated by stress-specific responses at different physiological and molecular levels. Stress-tolerant genotypes are a result of differential expression of unique proteins to protect essential plant functions when the plants are exposed to stress conditions. Understanding how plants respond while confronting abiotic stress is not only pivotal to discovery of stress response mechanisms, but has the potential to yield novel approaches for producing stress-resistant crops. Developing plant adaptation strategies to abiotic stresses is dependent on the discovery of these unique and shared stress-response mechanisms at different molecular levels in plants. Quantitative proteomics enables us to identify stress-specific proteins, which can then be used in determining the mechanisms underlying protein and gene targeting and trafficking, and the functions of specific cellular compartments in those processes.

### 3.1 Mild drought stressed Nipponbare plants respond differently to moderate and severe drought stressed plants

Nipponbare is known to have little tolerance for water deficits, so different levels of mild, moderate, and severe drought stress treatments were applied in this study to analyze the changes occurring in the plant proteome as a response. Increasing the water deficiency level from mild to severe drought showed that the number of DEPs increased significantly (Table 2). Moreover, a greater percentage of overlap between DEPs in moderate and severe drought stress illustrated that plants sensed and responded to these two drought levels similarly at the proteome level, and responded more dynamically than to mild drought levels.

Comparing the molecular functions involved in plant response to different drought stress levels, in both the unique and shared responses, revealed similar mechanisms and function alteration under all drought levels. For example, photosynthesis, transport, and translation were three major molecular functions that were changed in abundance the most under all three drought stress levels. Photosynthesis is one of the prime metabolic processes in plants affected by drought stress, and stress tolerance mechanisms in plants help them to maintain their photosynthetic status. Recent studies have shown a correlation between photosynthesis and transport functions, with water deficiency causing disruption of electron transport chain and subsequent inactivation of PSII [18, 19]. During water stress, plant cells must shift from growth to survival mode and alter metabolism towards functions critical for viability. For example, instead of shipping acetyl units to the cytosol, there is a greater requirement for acetyl-CoA to be oxidized in the mitochondria for ATP synthesis, so lower levels of nucleocytosolic acetyl-CoA limits growth-related processes [20]. This agrees with our observed decreased abundance of putative acyl-CoA dehydrogenase under the severe drought stress treatment, and decreased abundance of proteins in the growth functional category under all drought stress levels in plants.

Not all shared proteins were expressed similarly between different drought levels, but C2H2-type domain-containing protein and putative glycine-rich protein were two proteins with coordinated expression between mild and more severe drought levels. Glycine-rich protein (GRP) superfamily members are known to be involved in cellular stress responses and signalling in plants [21], although there are contrasting findings regarding changes in their abundance in response to drought stress. According to Shim et al, however, glycine-rich protein 3 functions as a positive regulator in drought tolerance in rice plants [22]. Similarly, C2H2-type domain-containing protein is a transcription factor that plays a critical role in the regulation of cellular and physical changes in response to environmental stresses in plants. Overexpression of a gene encoding a zinc finger protein regulates rice plant responses to drought stress by decreasing ROS [23, 24], accumulating free proline, and improving antioxidant enzyme activity [25]. Thus, our results support the observation that the Nipponbare genotype is resistant to mild drought level but threatened under stronger drought conditions. In addition, many molecular functions altered under drought stress responded to multi-level stress in different ways and at different levels.

### 3.2 The role of aquaporins in plant response to abiotic stresses

The transport of water across biological membranes in plant cells happens through specialized pores which enable them to transport water into and out of the cells along a water potential gradient, and these pores are composed predominantly of aquaporins [26], which include seven subfamilies categorized according to their intracellular locations and sequence similarities [27]. The plasma membrane intrinsic proteins (PIPs) and tonoplast intrinsic proteins (TIPs) were changed in abundance under individual abiotic stress in this study. For example, aquaporin PIP2-7 increased in abundance under drought stress and decreased in abundance in response to salinity, which is in agreement with the finding that heat and salt stress are known to commonly affect the transport and compartmentation of ions in plants [1]. Aquaporin TIP1-2 was decreased in abundance under temperature stress, which contrasted with what was observed for aquaporin PIP2-1, which showed an increase in abundance under the same stress condition. Several previous investigations have shown that PIPs function as transporters of water, glycerol, H2O2, carbon dioxide, and urea, and are also involved in abiotic stress responses. Moreover, several studies have demonstrated differential abundance of PIPs and TIPs family members in response to salt, drought, or cold stresses [28, 29]. Our results showed that different aquaporins were altered in abundance in different directions in response to various abiotic stresses, which confirms their crucial role in stress response in plants.

### 3.3 Individual stresses trigger changes in abundance of proteins with specific functions

Photosynthetic inhibition is one of the primary detrimental effects of water stress, due to stomatal closure, and one of the mechanisms plants have developed to cope with drought stress is recovery of photosynthesis by accumulating drought-responsive proteins involved in photoreactions [30]. In our data, 11 out of 39 proteins increased in abundance only in response to drought stress in Nipponbare plants were related to photosynthetic functions, such as two chloroplastic proteins, photosystem I iron-sulfur centre protein and cytochrome b6. Conversely, two proteins known to be negatively regulated in response to water deprivation in plants, Zeaxanthin epoxidase and Dehydrin DHN1, were decreased in abundance in our results. This is functionally similar to the finding of inhibition of Zeaxanthin epoxidase activity under oxidative stress in Spinach plants [31].

The activity of V-ATPase proteins is mediated by ion transport in plant cells, which could be affected by ionic stress during salinity. V-ATPases perform an important proton pump function in plant cells, regulating homoeostasis of cytosolic pH and playing a role in transport processes of the secretory pathway. Manipulation of expression of V-ATPases has been proposed as a potential means of improving crop yield while also improving stress resistance [32]. Moreover, the 14-3-3 protein family has been previously identified as a regulator of signaling pathways interacting positively with plasma membrane ion transporters [33]. This is consistent with our finding that two proteins, 14-3-3-like protein GF14-C and 14-3-3-like protein GF14-F, as well as two V-ATPase proteins, were reduced in abundance in response to salinity stress. Additionally, SGT1 (a ubiquitin ligase homolog), was increased in abundance in Nipponbare plants only after exposure to temperature stress. This is interesting, because a previous report showed that SGT1 plays an important role in plant response to a range of pathogenic infections, suggesting it is a multifunctional protein [34].

### 3.4 Delineation of protein cellular localization as stress-related factor

Protein biological functions and cellular localizations both play a pivotal role in determining stress-related protein accumulation [10]. Stress-sensing is thought to happen in the cell wall or cell membrane and is then relayed to various subcellular locations including the nucleus, cytosol, mitochondria, vacuole, chloroplast, and endoplasmic reticulum [7]. Furthermore, various cellular structures may be involved in abiotic stresses, both with common responses (osmotic and oxidative stresses) and stress-specific responses (ionic stress) [35].

The cytosol was the only organelle in our study where proteins responded to all individual abiotic stresses both positively and negatively at the proteome level. RNA-binding protein and phosphoglucomutase protein were two proteins that increased in abundance in common under all abiotic stresses, although their biological functions are quite different. RNA-binding proteins are likely to regulate more than 60% of the plant transcriptome [36], and overexpression of RNA-binding proteins strongly induces tolerance of Arabidopsis plants against environmental stresses [37]. Cytosolic phosphoglucomutase is essential for sucrose formation in plant cells, playing a crucial role in carbohydrate metabolism mechanisms in plants [38]. Another cytosol-localized protein was peptidyl-prolyl cis-trans isomerase, which is involved in a variety of cellular processes including protein folding. The positive role of the peptidyl-prolyl cis-trans isomerase in plant acclimation to temperature and salinity tolerance has been previously observed, however, its specific function under drought stress remains unclear [39].

The nucleus is a site where stress signals are transformed into gene expression, and any imbalance in cellular conditions is sensed by receptors at the plasma membrane, inducing signaling pathways which transfer stress signals to the nucleus and leading to changes in gene expression, signal transduction, and abundance of chaperones, ROS scavenging enzymes and regulatory proteins [10]. All abiotic stresses in our study affected the nucleus proteome negatively by decreasing the abundance of proteins involved with gene expression, signal transduction, and metabolic processes. Interestingly, about 40% of the proteins commonly decreased in abundance under all abiotic stresses were localized in the nucleus, including transcription factor proteins, calcium-dependent protein kinase 8, and ribokinase. Protein kinases are major players in various signal transduction pathways in plants, including many that are related to abiotic stresses [40]. Calcium-dependent protein kinase was one of the kinase protein family members in our study which showed a decrease in abundance under all stress treatments. This protein performs a signaling function in all cellular organelles including the nucleus, cytosol, and plasma membrane. Studies have revealed that temperature stress can change the fluidity of cellular phospholipid membranes by affecting kinase proteins [41], in agreement with the finding that salt stress effects kinase proteins acting in the plasma membrane of rice root cells [42].

## 4. Conclusion

In this experiment, we analyzed the common and specific proteome responses to stress of Nipponbare rice plants exposed to three major individual abiotic stresses. This enabled elucidation of the protein level responses that occur under various abiotic stress conditions, and their physiological significance. Both unique and common rice proteome responses were identified based on the severity and nature of the stresses involved. For example, severe and moderate drought stress levels induced similar changes in protein abundance in plants, in contrast to mild drought stress.

Comparing the proteome alteration between diverse abiotic stresses illustrated that the most common rice proteome responses identified under all abiotic stresses include substantial changes in the photosynthesis apparatus, protein transport, translation, redox homeostasis, and detoxification/antioxidation pathways. These all occurred in various cellular organelles including the nucleus, chloroplast, and plasma membrane. One family of proteins which clearly plays an important role in stress response is the aquaporins. In this study, a number of different aquaporins were increased or decreased in abundance in response to either specific stresses or various stress combinations.

Unique and shared proteome responses of plant leaf tissues to abiotic stresses can lead to finding both stress-specific signals and common responses. Identifying which proteins are differentially expressed, in which organelles, under which stress, helps to provide a more complete picture of how the plant cells are responding to stress at the molecular level. Therefore, subsequent functional characterization of key stress-tolerance proteins such as HSPs, aquaporins and kinases identified in this study could provide quantitative biomarkers of stress responses. Enhancing our understanding of stress signalling and responses will increase our ability to improve stress tolerance in crops, which is necessary in order to achieve agricultural sustainability and food security for a growing world population.

## 5. Materials and Methods

### 5.1 Plant growth and stress treatments

Five seeds of Nipponbare rice (Oryza sativa) were sown in the same pots (30 cm deep and 10 cm in diameter) filled with 700g soil weighed for each pot. There were 18 pots in total, comprising 3 replicates each of control, mild drought stress, moderate drought stress, severe drought stress, salt stress, and temperature stress. Plants were fertilized twice, the first time simultaneously with soil filling and again 2 weeks after sowing. Prior to sowing, seeds were sterilized in four steps; washing in 70% ethanol for 20 minutes, water for 1 minute, 50% bleach solution for 30 minutes, and final washing in water for 5 minutes. Plants were grown in a controlled greenhouse condition with temperature set to 28/22°C (day/night) and a 12-h photoperiod. Light intensity exceeded 700 μmolm^-2^s^-1^ throughout.

After 4 weeks of growth in optimal well-watered conditions, the seedling plants were subjected separately to a particular abiotic stress including: watering reduction to 30%, 50%, and 70% of soil field capacity (FC) as three levels of drought stress, 50mM NaCl concentration as salt stress, and 33/18°C day/night temperature regime as temperature stress. Leaf tissue samples were collected after 0 day (Control) and 6 days (Stress) and frozen immediately in liquid nitrogen. For further proteome analysis, Samples were placed in 2 ml centrifuge tubes and ground finely using a Qiagen Retsch 12090 TissueLyser II, five Zironox beads (2.8-3.3 mm,) and liquid nitrogen.

### 5.2 Protein extraction and assay

50 mg of leaf powder was suspended in 1.5 ml of 10% trichloroacetic acid in acetone, 0.07%β-mercaptoethanol, and incubated at −20°C for 45 minutes. The extract was centrifuged for 15 minutes at 16,000 ×g at 4°C, and the pellet was collected and washed with 1.5 ml of 100% acetone followed by centrifugation for 15 minutes at 16,000 ×g at 4°C. The acetone washing step was repeated three times for the complete removal of pigments, lipids, and other lipophilic molecules. The colorless resulting pellet was lyophilized in a vacuum centrifuge for 5 minutes. Then, 400 μl of 2% SDS in 50 mM Tris-HCl (pH 8.8) was used to resuspend the pellet. After vortexing for 2 hours, the pellet was removed and the supernatant was kept for reduction and alkylation. The samples were reduced by adding 1M Dithiothreitol (DTT) to reach a final concentration of 10mM and incubated for 1 hour at 37°C, followed by alkylation with 20 mM Iodoacetamide (IAA) for 45 minutes in the dark at room temperature.

Samples were then methanol-chloroform precipitated. 300 μl of protein solution was mixed with 800 μl of methanol and 200 μl of chloroform. 500 μl of water was added and vortexed and the mixture was centrifuged at 6,000 ×g for 2 minutes. After removing the upper phase, 600 μl of methanol was added to the mixture and centrifuged at 6,000 ×g for 2 minutes, and the supernatant was then removed. The pellet was air-dried and solubilized in 80 μl of 8 M urea in 100 mM Tris-HCl buffer (pH 8.8). The concentration of protein in the solution was measured by bicinchoninic acid (BCA) assay kit (Thermo Fisher Scientific, San Jose, USA).

### 5.3 Trypsin in-solution digestion and peptide extraction

200 μg aliquots of protein were used for digestion and peptide extraction. Samples were first diluted five-fold with 100 mM Tris-HCl buffer (pH8.8), then trypsin was added (1:50 enzyme: protein) and incubated overnight at 37°C. The reaction was stopped by adding trifluoroacetic acid (TFA) to reach a final concentration of 1%. Peptide samples were desalted using stage-tips (SDB-RPS, 3M, Saint Paul, USA). Samples were spun and loaded in stage tips (4 punches of SDB membrane used in 200 μl pipette tips), held by adaptors in 2 ml tubes, and centrifuged with maximum speed of 2500 rpm to pass the sample through the membrane. The tips were washed two times with 200 μl of 0.2% TFA and peptides were eluted by addition of 200 μl of 80% Acetonitrile (ACN) and 5% NH4OH solution. The peptide concentration was measured using a micro BCA kit (Thermo Fisher Scientific, San Jose, USA). Peptides were subsequently fractionated using high pH reversed-phase peptide fractionation kit (Thermo Fisher Scientific, San Jose, USA) and pooled into 8 fractions [43]. The fractions were dried and reconstituted in 1% formic acid.

### 5.4 Nano LC-MS/MS

Peptides were analyzed by nanoflow LC-MS/MS using a Q Exactive Orbitrap mass spectrometer coupled to an EASY-nLC1000 nano-flow HPLC system (Thermo Fisher Scientific, San Jose, CA, USA). Reversed-phase columns of 75 μm internal diameter were packed in-house to 15 cm length with ES-C18 Halo, 2.7 μm, 160 Å, (Advanced Materials Technology, Wilmington, DE, USA). Peptides were eluted from the column for 60 min, starting with 100% buffer A (0.1% formic acid), using a linear solvent gradient with steps from 2 to 30% of buffer B (99.9% (v/v) ACN, 0.1% (v/v) formic acid) for 50 min and 30 to 85% of buffer B for 10 min. One full MS scan over the scan range of 350 to 1850 m/z was acquired in the Orbitrap at a resolution of 70,000 after accumulation to automated gain control (AGC) target value of 1 × 10^7^. MS/MS fragmentation spectra were acquired for the 10 most intense ions. The maximum injection time was set to 60 ms and higher-energy collisional dissociation fragmentation was performed at 27% normalized collision energy, with selected ions dynamically excluded for 20s.

### 5.5 Data processing

The raw data were converted to mzXML format and searched against available O. Sativa protein sequences in UniProt (48904 sequence entries, downloaded April 2020) using the X!Tandem algorithm operating within the global proteome machine software (GPM, version 3.0, www.thegpm.org). Peptide to spectrum matching parameters included a mass tolerance of ± 10 ppm for the parent ion and 0.4 Da for the fragment ion, log (e) values less than −1 for peptides and proteins, tolerance of one missed tryptic cleavage, complete modification of cysteine by carbamidomethylation, and potential modification of methionine by oxidation. As a result, 18 merged output files containing peptides and proteins identified from 8 fractions from each sample were generated. All raw data have been submitted to the PRIDE data repository, and will be available with project identifier PXD037280.

### 5.6 Analysis of quantitative proteomics data outputs

To create a single merged list of reproducibly identified proteins, three biological replicates were combined into a merged list using Scrappy [44] and false discovery rates (FDR) were calculated at the protein and peptide levels for each. The criteria for a reproducibly identified protein to be retained in the merged data set were that it was identified in all three biological replicates of at least one condition, and the total spectral count across the three replicates was at least six peptides [45]. Protein abundances were calculated using Normalized Spectral Abundance Factors (NSAF) with the addition of a spectral fraction of 0.5 to all spectral counts to compensate for null values and allow log transformation for additional statistical analyses [44]. Pairwise comparisons of the stress versus control condition in different treatments were performed using student t-tests of the log-transformed NSAF data. Proteins with a p-value less than 0.05 were considered as statistically significantly differentially expressed [46]. Fold changes were calculated as a ratio of the averaged log NSAF value of proteins present under individual abiotic stresses compared to those under control conditions.

### 5.7 Parallel Reaction Monitoring Analysis

Parallel Reaction Monitoring (PRM) analysis was used for the validation of the label-free shotgun proteomics results, to measure quantitative changes in specific proteins in Nipponbare plant samples. Skyline (ver. 21.2.0.568) was used to create an inclusion list of unique peptides, for each selected protein, with their mass to charge ratio [47]. The obtained PRM data were imported into Skyline and subject to quality control analysis. The sum of areas of all transition peaks for each peptide were collected and log2-transformed, and a Student t-test analysis was used to compare protein abundance between stress and control conditions.

## 6. Acknowledgements

This work was supported by Australian Research Council Discovery Project DP190103140 and Macquarie University, and aspects of this research were conducted at the Australian Proteome Analysis Facility. Supplementary data are available on request.

## Notes

### Competing Interest Statement

The authors have declared no competing interest.

